# DeepRNAi: Predicting on- and off-target siRNA activity and establishing a benchmark

**DOI:** 10.64898/2026.11.12.766311

**Authors:** Nicolas Y. Masse, Tammy T.Y. Lau, Jeff Wintersinger, Mykhaylo Slobodyanyuk, Roujia Li, Vivian S. Chu, Helen Zhu, Bhargav Kanuparthi, Jonathan La, Brodie Fischbacher, Krista Arseneault, Sydney Johnson, Wang-Ting Lu, Noah Lorincz-Comi, Emma Bohn, Christopher B. Cole, Gregory R. Hoffman, Nathan H. Lazar, Scott D. Findlay, Albi Celaj, Brendan J. Frey

**Affiliations:** Deep Genomics Inc., Toronto, Ontario, Canada; Deep Genomics Inc., Cambridge, Massachusetts, USA

**Keywords:** siRNA, RNAi, RNA interference, microRNA, oligo therapeutics, genetic medicine, off-target prediction, specificity, transcriptome, RNA-seq, biological foundation model, BioFM, virtual screening

## Abstract

Small interfering RNAs (siRNAs) silence target genes through RISC-mediated hybridization and cleavage, but can silence unintended transcripts and cause toxicity or complicate regulatory approval. In drug development, empirical assessment by qPCR and RNA-seq is slow, forces early candidate prioritization, and is limited by available species- and genome-appropriate cell models. While extensive empirical screening remains standard practice, there is a significant opportunity for modern AI methods to change this paradigm. We introduce DeepRNAi, a deep learning model that accurately predicts on- and off-target knockdown from siRNA and mRNA sequence. Trained on 16 million siRNA-transcript pairs from 2,031 siRNAs designed to interrogate known siRNA failure modes, and incorporating transcript context from biological foundation models, DeepRNAi is the first *in silico* method that approaches experimental accuracy. It generalizes across cell models, transcripts, genomes and species, and in a small study distinguishes approved therapeutics from hepatotoxic candidates. The value of DeepRNAi for prioritizing better candidates up front is supported by a genome-wide survey of 24.4 million siRNAs targeting 7,946 genes, which indicates that 90.1% of the 100 most potent siRNAs have at least some off-target concerns. To support the community, we are publishing a training and benchmark dataset containing 1.84 million siRNA-transcript pairs from 240 siRNAs.

## Introduction

RNA interference (RNAi) converts short RNA duplexes into programmable regulators of gene expression. The guide strand of a small interfering RNA (siRNA) is loaded into an Argonaute-containing RNA-induced silencing complex (RISC), which recognizes complementary messenger RNAs and promotes their cleavage or destabilization.^1–3^ Improvements in chemical stabilization, conjugate-mediated delivery and clinical development have turned this mechanism into a validated therapeutic modality.^4^ By acting upstream of protein production, siRNAs can address targets that other modalities cannot, including proteins that lack a druggable binding pocket and intracellular targets inaccessible to antibodies.^5^ They can be remarkably potent and long lasting, with some approved treatments requiring only semi-annual maintenance dosing.^6^ siRNA drug design is particularly well suited to computational modeling since each candidate is specified by a discrete sequence and a defined set of chemical modifications. However, off-target interactions can lead to unintended knockdown of other transcripts that may cause toxicity and complicate regulatory approval. While qPCR and RNA-seq can assess a subset of possible candidates in cell models, doing so can introduce delays, prematurely narrow the candidate pool to accommodate experimental capacity, and lead to incorrect prioritization due to differences in cell contexts.^7–9^ Computational methods can be much faster, keep the funnel broader, and generalize across cell models, genomes and species to help guide therapeutic program decisions such as animal model selection. However, until now, these methods have shown limited utility, so extensive screening, especially in human cell lines to assess off-target burden, remains standard practice.^5,10–12^

The two dominant classes of hybridization-dependent off-target activity are due to near-perfect matches (NPMs), which can support extensive guide-target pairing and cleavage-like repression, and seed matches, which generate microRNA-like repression through pairing between guide positions 2–8 of the guide strand and the mRNA sequence, especially in the 3′ untranslated region (3′ UTR).^10,13,14^ Aggregate transcriptome-wide seed-mediated off-target activity has been associated with hepatotoxicity in preclinical studies and has even led to setbacks in clinical programs.^7,15^ Hybridization-dependent off-target liabilities also create development and regulatory burdens, as FDA draft guidance recommends their assessment and experimental follow-up.^12,16^ While chemical modifications can be used to reduce sequence-mediated liabilities for siRNA candidates,^13,17^ optimizing these modifications introduces additional delays and design complexity. Modification is also typically applied only after the pool has been narrowed by initial candidate selection, so candidates that would have performed best once modified may already have been discarded. It is therefore preferable to evaluate a much broader set of sequences for potency and specificity early in the design process.

Despite the importance of designing sequences with low off-target burden, no established predictive model has been broadly validated for prospectively predicting the transcriptome-wide off-target effects of therapeutic siRNAs before experimental testing. Most siRNA-specific predictive methods instead focus only on on-target potency,^18–20^ whereas bioinformatic off-target approaches primarily identify sequence matches,^21–23^ of which only 4% to 8% will validate as functional off-target sites. The closest precedent for transcriptome-wide prediction of seed-mediated repression comes from miRNA work, most notably TargetScan,^24,25^ which predicts repression using features such as the extent of seed pairing, seed-pairing stability, target-site context, and binding affinity. Our previous work on the REPRESS model for predicting cell-type-specific microRNA binding and mRNA degradation from transcript sequence is relevant here, but it does not alone predict the response to an exogenous siRNA.^9^ There is therefore a critical need for models trained directly on siRNA perturbation data that can predict transcriptome-wide responses to therapeutic siRNAs and derisk prospective designs before experimental testing.

Predicting siRNA-mediated repression is challenging because activity can depend on features operating at multiple scales, including siRNA-specific features such as base and motif composition, duplex free energy and seed region characteristics^26–29^; local siRNA-mRNA binding site features such as siRNA-mRNA interaction thermodynamics, target-site accessibility,^30^ position within the transcript,^31^ and local secondary structure^32^; and mRNA-level properties such as stability and expression level.^8,24^ While narrowly focused models have been developed for on-target prediction^18,19,33^ and certain kinds of off-target prediction,^21,22,34–36^ to date no model has combined siRNA, local binding-site and transcript-level features to predict transcriptome-wide off-target repression at a performance level meaningful for therapeutic design.

We introduce DeepRNAi, a deep learning model that predicts siRNA-induced on- and off-target repression across the transcriptome, together with a benchmark dataset from 240 siRNAs comprising 1.8 million siRNA-mRNA interaction signals. Three elements are central to the approach: a fit-for-purpose differential expression dataset enriched for sequence-mediated interactions; preprocessing that focuses learning on direct hybridization-mediated knockdown; and an architecture integrating multi-scale context from RNA biological foundation model (BioFM) embeddings and experimental covariates. DeepRNAi substantially outperforms prior approaches including TargetScan, with predictive accuracy approaching the reproducibility of RNA-seq measurements across cell lines and across independent screens in the same cell line. It generalizes to transcripts, genomes and species not seen during training, and its predictions reflect interpretable biology. For example, DeepRNAi predicted reduced activity for siRNAs targeting a 3′ UTR subregion of the *PLG* gene subject to genetically driven alternative polyadenylation, consistent with experimental data.

To assess how off-target liabilities may impact siRNA drug development generally, we used DeepRNAi to evaluate 24.4 million siRNAs targeting 7,946 human protein-coding genes and found that acceptably low off-target burden is rare among potent siRNAs, underscoring the need for model-guided design to identify high-specificity candidates.

Together, these results demonstrate that DeepRNAi is a frontier model for siRNA design that provides substantial gains both on technical metrics and on metrics reflecting real impact on drug discovery. DeepRNAi accurately predicts siRNA-induced, transcriptome-wide changes in gene expression, enabling prospective prediction of siRNA specificity, upfront prioritization of potent candidates with minimal off-target burden, and rapid, large-scale exploration of sequence design space far beyond what can be tested experimentally.

## Results

### DeepRNAi accurately predicts siRNA effects *in vitro*

To train DeepRNAi, we generated a fit-for-purpose differential expression RNA-seq dataset in HepG2 and Huh7 cells consisting of 2,271 siRNAs, targeting 71 genes, screened at 10 nM for 48 hours. To enrich the training set for a substantial number of siRNA-mRNA interactions, we intentionally designed siRNAs enriched for NPMs to transcripts expressed in these cell lines (7,946 genes). This dataset contains more than 17 million siRNA-mRNA pairs, including approximately 388,000 pairs with observed log2 fold change (log_2_FC) < −0.5 and adjusted p-value < 0.05. We used 2,031 siRNAs targeting 62 genes for training, and constructed a held-out evaluation set of 240 siRNAs targeting 9 genes, in which neither the guide sequences, nor any other guides targeting the same target transcripts, or any paralogs, were in the training set. This target-transcript holdout is intentionally harder than a random siRNA split, and evaluates the ability for the model to enable “zero-shot” design for an arbitrary new therapeutic program without requiring any target-specific screening data for training. The held-out evaluation set comprises the public benchmark dataset as described below.

DeepRNAi models the response of any candidate siRNA-mRNA pair by jointly taking into account the siRNA guide sequence (the 5′ most 21mer), the on- or off-target mRNA sequence, and categorical variables for cell line, siRNA format, and chemistry (**Figure 1a**). These representations are integrated with single-nucleotide-resolution embeddings from a BioFM to predict the degree of hybridization-dependent knockdown (log_2_FC) for any siRNA-mRNA pair. To focus model training on direct, RISC-mediated knockdown, we developed an unsupervised algorithm that reduces estimated indirect effects, noise and other confounding factors to enrich for direct, sequence specific and hybridization-dependent signals, and trained DeepRNAi on this de-noised signal. To assess model accuracy, we compared predicted and observed log_2_FC values on a held-out test set after centering each transcript’s observed log_2_FC values using the mean calculated across all training-set siRNAs. This reduces the contribution of shared/consistent transcript responses to measured performance without using test set information (**Methods**).

**Figure 1:**
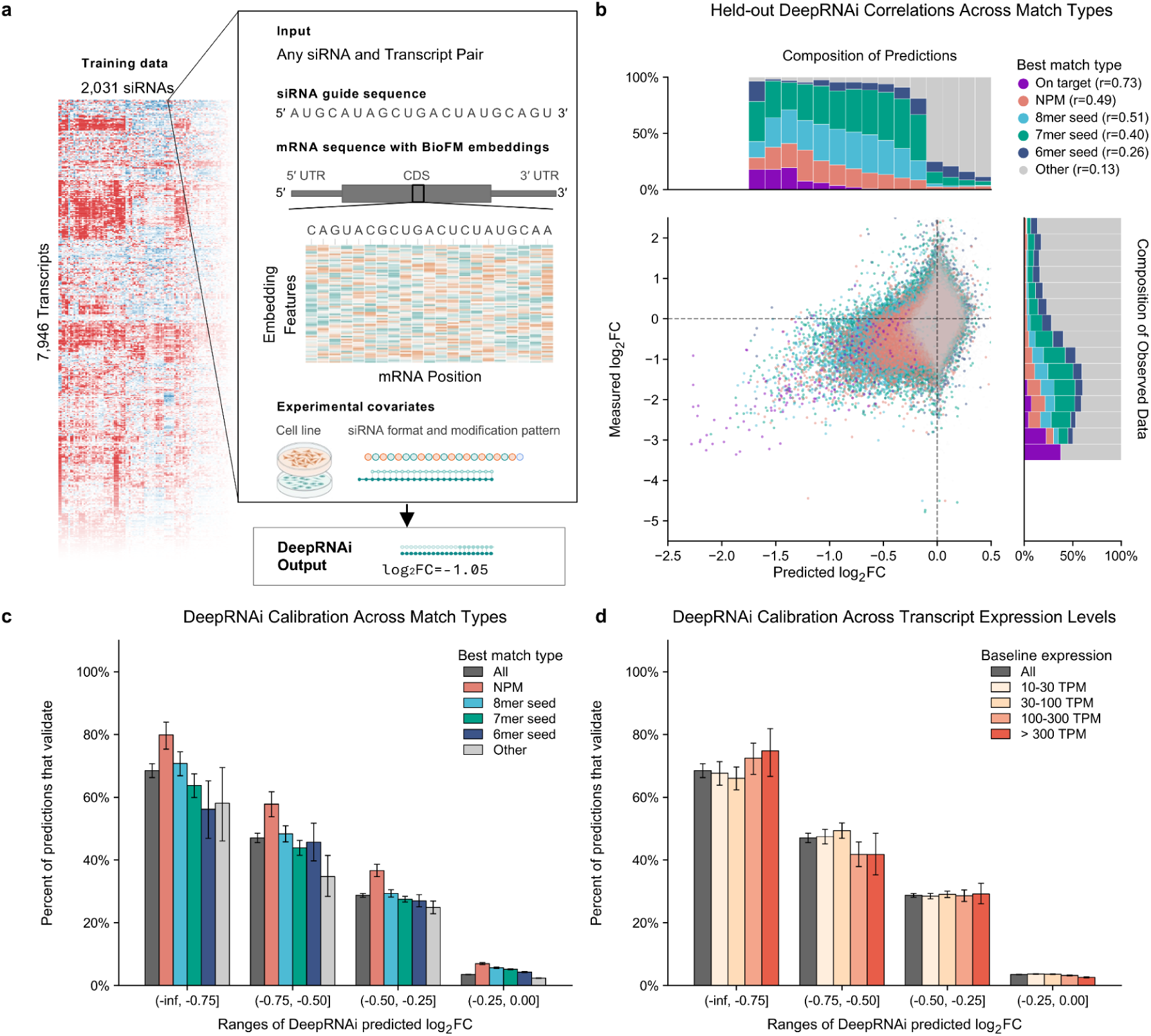
DeepRNAi is a state of the art predictor for RNAi off-target effects. **a)** Representation of the training data, inputs, and output predictions for the DeepRNAi model. **b)** Scatter plot of predicted vs observed log2 fold-change (log_2_FC) values for the DeepRNAi model using the held-out test set. Each point represents a single siRNA-mRNA pair and is colored by the highest-ranking type of match between the siRNA and mRNA (NPM: near-perfect match). Distributions of match types are shown in margins and Pearson *r* correlation coefficients in the legend. **c)** Precision of DeepRNAi predictions, binned by predicted magnitude of knockdown, stratified by degree of complementarity. A prediction for an siRNA-mRNA pair is considered validated if the observed log_2_FC < -0.5 with adjusted *p* < 0.05. Error bars show 95% Clopper-Pearson binomial confidence intervals. **d)** Same as (c) except stratified by expression of the off-target transcript.

We assessed DeepRNAi’s performance across several classes of sequence complementarity: intended on-targets (perfect complementarity to the siRNA guide); near-perfect matches (NPMs) based on positions 2–16 of the siRNA guide strand with a maximum edit distance of two; and seed matches to the 3′ UTR of the transcript. Seed matches were further classified into established seed-match subtypes (6mer, 7mer-A1, 7mer-m8, and 8mer)^24^ (**Methods**). Annotations were mutually exclusive, with each siRNA-mRNA pair assigned its highest-ranking match type according to the following hierarchy: on-target > NPM > 8mer > 7mer-m8 > 7mer-A1 > 6mer.

Correlations were highest for intended on-target transcripts (*r* = 0.73, *n* = 229), followed by NPMs (*r* = 0.49, *n* = 51,345), and seed matches in the 3′ UTR, which ranged from *r* = 0.51 (*n* = 56,642) for 8-mers to *r* = 0.26 (*n* = 160,837) for 6-mers (**Figure 1b**). Predicted effect size was also well calibrated, with increasingly negative predictions yielding progressively larger proportions of validated hits (log_2_FC < -0.5 and adjusted *p* < 0.05), regardless of sequence complementarity and transcript expression levels (**Figures 1c,d**).

To benchmark DeepRNAi against an established model and a conventional machine learning baseline, we evaluated its performance against TargetScan Human 7.0^24^ and a Random Forest (RF) model trained on the same proprietary Deep Genomics dataset (**Methods**). We compared models across siRNA-mRNA pairs with perfect complementarity (i.e. intended on-targets), NPMs and seed matches. Since TargetScan only generates predictions for seed matches, we only show its accuracy for this class of siRNA-mRNA pairs.

For intended on-targets, DeepRNAi significantly outperforms the RF model (*p* = 0.021, paired bootstrap, **Methods**) (**Figure 2a** bar plot, **Table S1**). We were also interested in whether BioFM embeddings provide DeepRNAi information that is being used to improve model accuracy. Thus, we retrained and evaluated a version of the model without the BioFM embeddings. On-target accuracy for this model was significantly below that of the full model (*p* = 0.004), confirming that BioFMs provide useful representations for on-target prediction accuracy.

**Figure 2:**
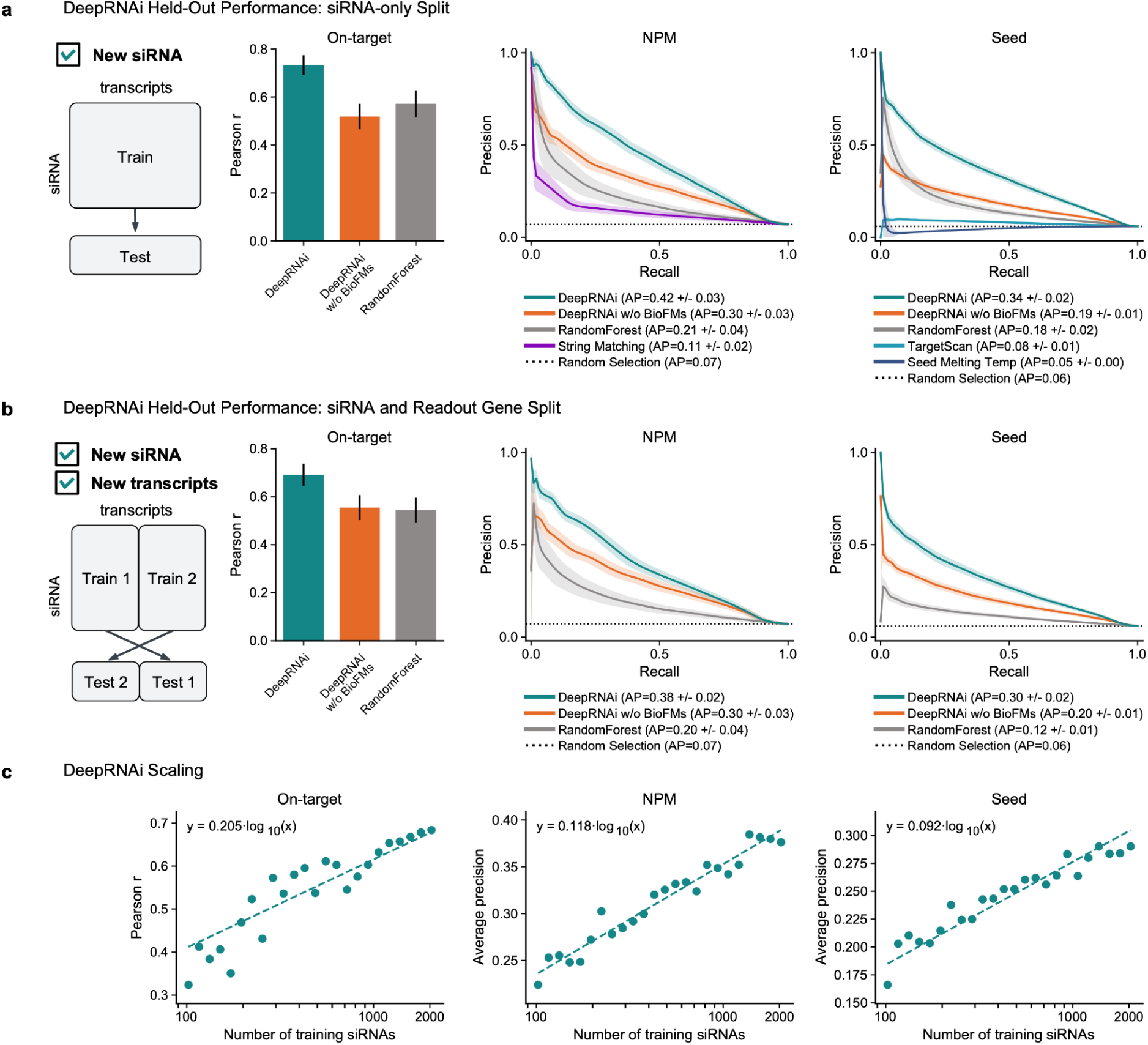
DeepRNAi performance on challenging *in vitro* tasks. **a)** Pearson correlation for on-target knockdown and precision-recall curves (PRCs) with average precision (AP) for classifying NPM and seed match knockdown (log_2_FC < -0.5 and adjusted *p* < 0.05) for test siRNAs targeting genes that were not seen in the training data. Metrics are shown for DeepRNAi, a version of DeepRNAi that was trained without BioFMs (DeepRNAi w/o BioFMs) and a random forest model (RF). A comparison to string matching is shown for NPMs and comparisons to both TargetScan and seed Tm are shown for seed matches. The error bars for the barplot and PRC curves represent ± one standard deviation (bootstrap). **b)** Performance comparisons as in (a) for models trained using only half of the transcript readout measurements, testing on the other half. **c)** Scaling plots showing Pearson correlation for on-target knockdown prediction and AP for classifying knockdown of NPM and seed match off-target siRNA-mRNA pairs for DeepRNAi-like models trained with variably sized datasets. Fit lines with zero intercept are annotated.

An important challenge in siRNA design is avoiding off-target NPMs or seed matches that produce meaningful transcript knockdown, given the abundance of bioinformatically plausible but non-functional binding events. To assess performance at this task, we generated precision-recall curves (PRCs) and calculated the average precision (AP) for all siRNA-mRNA pairs with either an NPM or seed match. A knockdown was deemed functional (positive) if observed log_2_FC < -0.5 and adjusted p-value < 0.05, and not functional (negative) if observed log_2_FC > -0.5 (regardless of p-value). PRCs show the performance of ranking positive and negative cases across all potential thresholds; in a perfect prediction, the plot would show a straight line across the top at a precision of 1.0 for all recall values. Similar to on-target accuracy, DeepRNAi significantly outperformed RF and string matching (**Methods**) on NPMs (*p* < 0.001) and significantly outperformed RF, TS and melting temperature (Tm) on seed matches (*p* < 0.001) (**Figure 2a** PRCs, **Table S1**). Also similar to the on-target results, DeepRNAi re-trained and evaluated without BioFMs substantially underperformed the full DeepRNAi model on both match types (*p* < 0.001), further confirming that BioFMs provide useful information DeepRNAi learned to leverage to improve predictive accuracy.

To evaluate model performance when predicting for unseen transcripts (e.g. in a new cell type or genome), we separated transcripts into two equal-sized disjoint sets and trained two separate models. Predictions from these models on each held-out transcript test set were then combined and evaluated as above (**Figure 2b**, **Table S2**). Similar to the results above, DeepRNAi significantly outperformed the RF model on intended on-targets (*p* = 0.03), NPMs (*p* = 0.002), and seed matches (*p* < 0.001). This test showed that DeepRNAi is able to successfully generalize to transcripts that it had not seen during training. In fact, its accuracy when trained on a subset of transcripts was not significantly different from the model trained on a dataset of equal size consisting of 50% of siRNAs and all transcripts (on-target: transcript split *r* = 0.69, 50% training data *r* = 0.70, *p* = 0.75, paired bootstrap; NPMs: transcript split AP = 0.38, 50% training data AP = 0.39, *p* = 0.45; seed matches: transcript split AP = 0.30, 50% training data AP = 0.31, *p* = 0.37) (**Figure S1**).

Scaling laws provide an important diagnostic for how deep learning methods perform with data and model complexity. In order to assess how model accuracy varies with the size of the training set, we retrained models on random subsets of siRNAs at 24 logarithmically spaced fractions ranging from 5% to 100% of our training dataset. Evaluating on the full test set we found that performance increased approximately log-linearly with data size for the three tasks described above, indicating a strong scaling law and suggesting that additional training data could further improve performance (**Figure 2c**).

### DeepRNAi accuracy approaches cross-cell-line reproducibility

We next compared DeepRNAi’s predictive performance with that obtained using observed responses to the same siRNAs in a different cell line. This comparison benchmarks model performance against the reproducibility of differential expression measurements across cellular contexts. We compared off-target accuracy on 11 siRNAs in our existing test dataset, screened independently in HepG2 and Huh7 cells (**Figure 3a**) and calculated the average precision using the observed response in one cell line as the target, and the observed response in the other cell line as the “prediction” (**Methods**). For both NPMs and seed matches, DeepRNAi model accuracy approaches that of RNA-seq reproducibility (**Table S3**), achieving approximately 74% (ranging from 63% to 80%) of the AP above chance baseline across both cell lines and match types.

**Figure 3:**
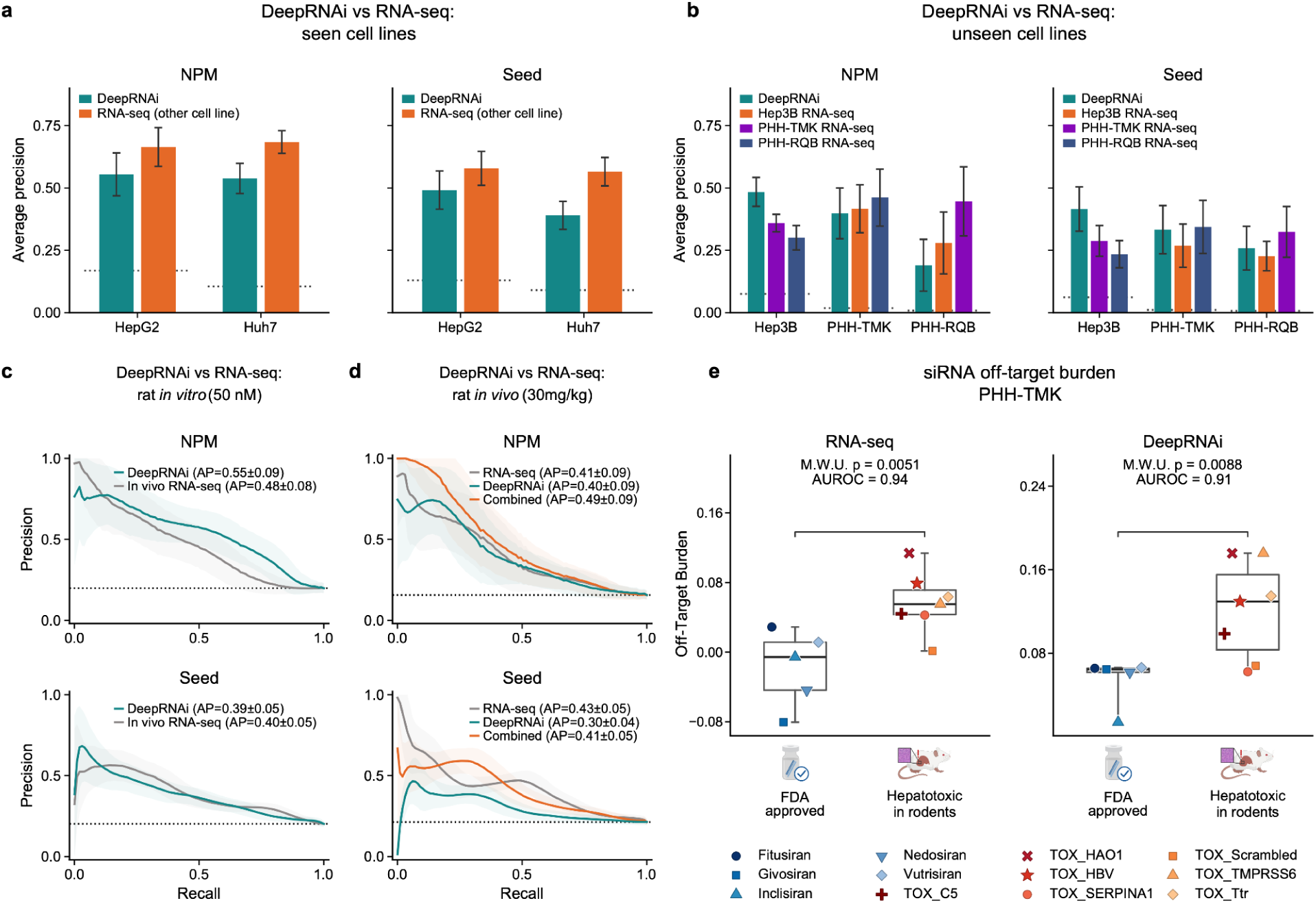
DeepRNAi generalizes to different cell lines, to different species (*in vitro* and *in vivo*), and to predicting toxicity liabilities. **a)** Barplots for DeepRNAi (cyan) against cross-cell line RNAseq reproducibility (orange) in which the observed values from one cell line are used to predict the observed values in the other. Left: NPMs, right: seed matches, the dashed lines show baseline positive rate. **b)** Same as a), except for three cell lines not found in the training data. **c)** Precision-Recall Curves (PRCs) and average precision (AP) of DeepRNAi predicted log_2_FC values for seed- and NPM-mediated expression changes in primary rat hepatocytes treated with two total siRNAs targeting two genes: *Hao1* and *Ttr*.^17^ Performance when using the *in vivo* RNA-seq data from d) as a predictor is shown for comparison. **d)** PRCs and AP for prediction of knockdown of off-target NPMs and seed matches in *in vivo* rat data (30 mg/kg dose *n* = 173 NPMs, 27 positive class, 146 negative class; *n* = 495 seed matches, 105 positive class, 390 negative class). DeepRNAi is compared with using *in vitro* rat hepatocyte log_2_FC data at 50nM to predict the *in vivo* responses. **e)** Box plots showing off-target burden between approved therapeutic siRNA (*n*=5) and siRNA believed to show rat toxicity in Janas et al.^7^ (*n*=7).Observed data is in PHH cells, off-target burden is defined as the difference in mean log_2_FC (restricted to log_2_FC < 0) between transcripts without NPM or seed matches and transcripts with matches. One-sided Mann-Whitney U test p-values and the Area Under the Receiver Operating Characteristic curve (AUROC) values are annotated.

We also tested whether DeepRNAi accuracy approaches that of RNA-seq derived differential expression for 15 siRNAs screened in the same cell line, but in different screens (**Methods**). Model accuracy was similar between DeepRNAi and RNA-seq for both NPMs and seed matches with DeepRNAi outperforming RNA-seq for NPMs and achieving 98% of the baseline-subtracted AP for seed matches (**Figure S2**, **Table S4**), further confirming that our model accuracy approaches that of experiment reproducibility.

### DeepRNAi’s predictions are accurate across cellular contexts, species and *in vivo* experiments

A persistent problem in the field is that predictions may be heavily confounded by training conditions that are unlike the conditions at application time. To address whether DeepRNAi overcomes this issue, we next asked whether DeepRNAi had learned the generalizable, hybridization-dependent effects of siRNA response rather than experiment-specific, cell-state-specific or otherwise indirect expression changes that might be only applicable to its training and evaluation datasets. We evaluated whether its predictions generalize across increasingly challenging settings: distinct human cell lines, an unseen species, and ultimately in an *in vivo* setting.

To assess generalization to novel human cell lines, we assessed model performance on data from 26 siRNAs screened in three novel cell lines not seen during training: Hep3B and Primary Human Hepatocyte (PHH) cells derived from two different donors. We repeated the same analysis as above where we compared model predictions to using differential expression data measured in different cell lines as a predictor. When comparing the baseline-subtracted AP across the three cell lines and the two match types (NPMs and seed matches), DeepRNAi predictions outperformed PHH cell types when predicting in Hep3B cells (144%-210%), performed similarly to Hep3B when predicting PHH (68%-124%), and performed slightly worse than using PHH from one donor to predict the other donor (42%-98%) (**Figure 3b**, **Table S5**). This suggests that DeepRNAi can generalize to novel types with accuracy approaching cross-cell line reproducibility.

Predicting off-target effects across species is helpful in minimizing hybridization-dependent toxicity risk in animal studies, interpreting any toxicity signals that are observed, and in selecting which animal models are ideal for a given lead candidate (e.g. rat versus mouse). Conversely, the absence of such signals can be misread as evidence that a molecule will be safe in humans. To address this, we tested whether DeepRNAi generalizes across species. DeepRNAi was trained exclusively on human HepG2 and Huh7 data, with no data from other species used for training or fine-tuning. Nevertheless, when applied to rat transcripts, its predictions agreed with NPM and seed-mediated expression changes measured in primary rat hepatocytes (**Methods**).^17^ For two siRNAs targeting *Hao1* and *Ttr* that were screened at 50 nM, DeepRNAi achieved an average precision of 0.55, compared with a baseline hit rate of 0.20, for classifying functional off-target NPMs. For seed matches, DeepRNAi achieved an average precision of 0.39, compared to a baseline hit rate of 0.20 (**Figure 3c**). These values are comparable to the performance of DeepRNAi on the tasks shown in **Figure 2**, suggesting minimal performance degradation when shifting to a new species context.

Having established that DeepRNAi generalizes across cell types and species *in vitro*, our next generalization test was to predict *in vivo* off-target effects in rat liver from the same study. Across 173 NPMs for the same *Hao1*- and *Ttr*-targeting siRNAs dosed at 30 mg/kg, DeepRNAi achieved an average precision of 0.40, compared with a baseline hit rate of 0.16, again reaching a similar performance to our *in vitro* test set (AP = 0.42 in **Figure 2a**) (**Figure 3d, Figure S3a)**. Strikingly, its performance was comparable to using experimental rat hepatocyte measurements from the same siRNAs to predict the *in vivo* NPM outcomes, which achieved an average precision of 0.41. Furthermore, when DeepRNAi’s predictions were used in combination with the rat hepatocyte measurements, this resulted in an average precision that exceeded both methods (AP = 0.49) (**Methods**), indicating both that DeepRNAi can be combined with noisy differential expression data from an *in vitro* setting to improve *in vivo* prediction, and that the gains are modest because DeepRNAi alone is able to predict most of the signal, without requiring *any* data for a given siRNA.

To summarize these results, although DeepRNAi was trained on human *in vitro* data, it generalizes zero-shot to predicting off-target effects in rat liver *in vivo* and matches the performance of experimental rat-hepatocyte measurements for this benchmark of classifying functional NPMs.

### DeepRNAi distinguishes approved therapeutics from hepatotoxic candidates

To explore its downstream utility for predicting clinical and late stage preclinical outcomes, we evaluated whether DeepRNAi predictions could accurately distinguish between FDA approved siRNAs and siRNAs that were found to be toxic in rodent studies due to off-target mediated effects.^7^ To that end we devised a simple metric for total “off-target burden” for a given siRNA: the difference in mean log_2_FC between a positive set of transcripts that contain an NPM or seed match and a negative set of all transcripts without NPM and seed matches (**Methods**). We computed this metric either using observed log_2_FC values from four different cell lines or using DeepRNAi predictions. We report both the one-sided Mann-Whitney U test p-value and the Area Under the Receiver Operating Characteristic curve (AUROC) which gives the probability of ranking a positive example (hepatotoxic, *n* = 7) over a negative example (FDA approved molecule, *n* = 5 available at the time of the study). For all four cell lines, we found a significant difference between the two groups for both the observed data (p-values from 0.001 to 0.037, AUROCs from 0.83 to 1.0) and the predictions (p-values from 0.009 to 0.015 and AUROCs from 0.89 to 0.91) (**Figure 3e, Figure S3b**). Although the small number of siRNAs warrants caution when interpreting these results, these findings suggest that DeepRNAi could help flag toxic candidates before advancement to *in vivo* studies.

### BioFMs improve DeepRNAi’s performance by predicting biology at scale

Early stages of siRNA drug discovery usually involve tiling against reference transcripts with all possible fully complementary guide strands of a given length. But this can lead to errors if the wrong transcript is chosen, annotations are incorrect, comprehensive annotations are not available, or RNA processing mechanisms such as polyadenylation and splicing are not accounted for. These issues are compounded across transcripts, cell models, animal species and individual genomes (**Figure 4a**). Here, we examine how BioFMs enable DeepRNAi to account for these factors to make predictions prospectively and at scale.

**Figure 4:**
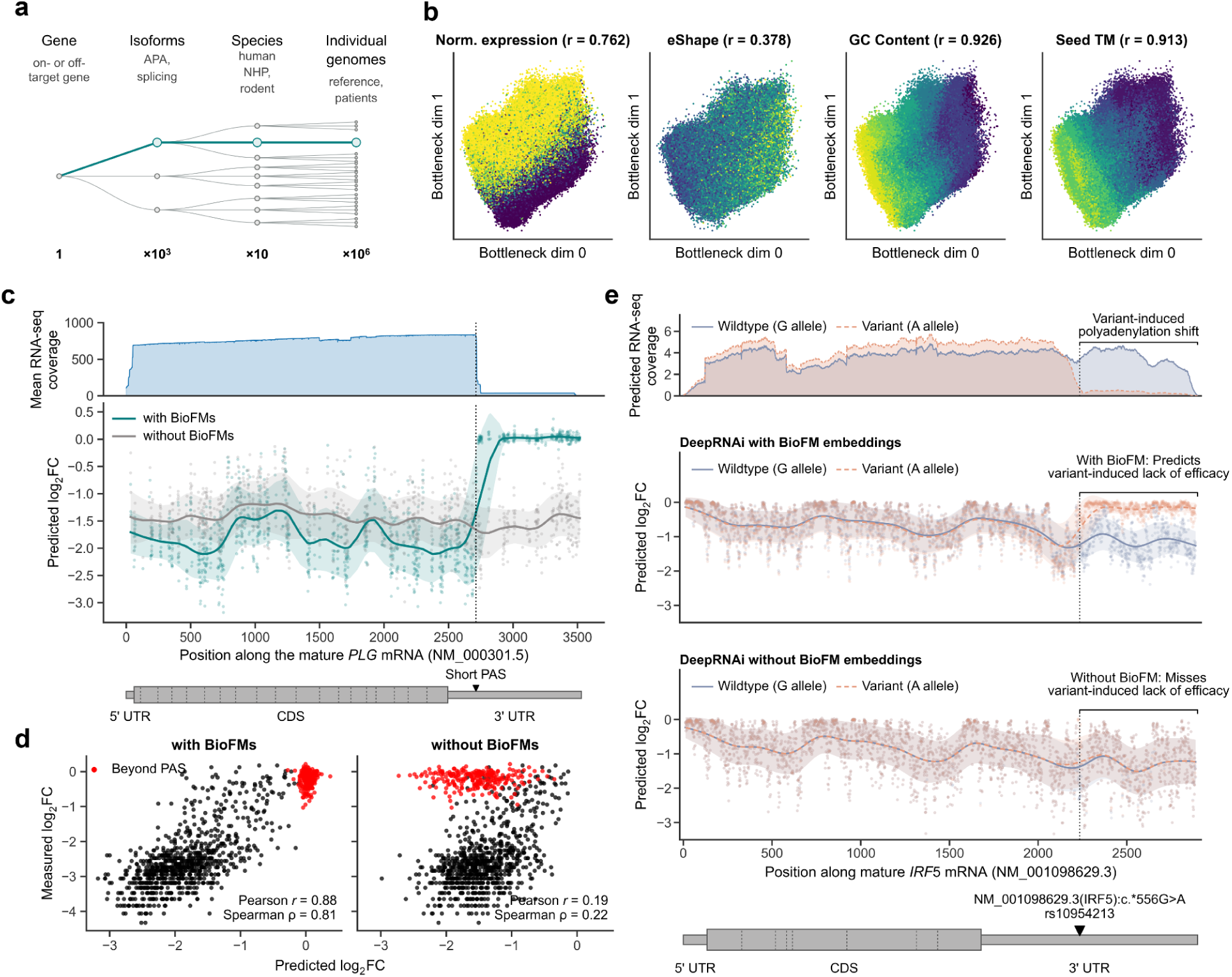
DeepRNAi encodes relevant genomic context. **a)** Schematic of the expansion of the number of contexts available and typically evaluated. **b)** 2D visualization of DeepRNAi’s internal representations. Attributes related to siRNA knockdown efficacy (normalized expression and eSHAPE accessibility of the mRNA at the putative target site, GC content and seed Tm of the guide) were read out from the activations in a late-stage DeepRNAi layer. Activations from this layer were first projected into 2D, and then projected onto the attribute predictions. Each dot is an siRNA and its color indicates the value of the attribute at that 2D location. The correlation between the predicted and actual attribute values is shown. **c)** Top: Read count coverage from long-read RNA-seq in HepG2 cells averaged over three replicates (**Methods**). Middle: DeepRNAi predicted on-target efficacy for siRNAs disclosed in a patent for *PLG* (PCT patent application WO2024077148A1, 2024) from models with and without BioFM embeddings. Bottom: Diagram of the NM_000301.5 *PLG* transcript, denoting the location of the proximal polyadenylation site (PAS), dotted vertical lines show exon boundaries **d)** Scatter plot, Pearson *r* and Spearman *ρ* of predicted and measured log_2_FC for siRNAs from c). siRNA targeting the 3′ UTR past the indicated PAS are shown in red. Left: predictions made with a DeepRNAi model trained with BioFM embeddings (Pearson correlation for points upstream of the short PAS: *r* = 0.68). Right: predictions made with a DeepRNAi model that was trained without BioFM embeddings (Pearson correlation for points upstream of the short PAS *r* = 0.52). **e)** Top: RNA-seq coverage predicted by Enigma^44^ for the NM_001098629.3 transcript of *IRF5* with (MUT) and without (WT) a single-nucleotide variant (chr7:128949373:G:A). Middle: DeepRNAi predicted on-target efficacy for siRNAs targeting NM_001098629.3 when the wild-type (WT) or mutant (MT) sequence is used as input to DeepRNAi with BioFM embeddings (center top) and without BioFM embeddings (center bottom). Bottom: Diagram of the NM_001098629.3 *IRF5* transcript, denoting the location of the variant, dotted vertical lines show exon boundaries.

First, we ‘opened up the black box’ and looked to see whether DeepRNAi has learned to account for previously studied biophysical attributes, such as RNA accessibility, seed Tm and GC content. We assessed the Pearson correlation between eSHAPE transcript accessibility measurements^37^ and the observed log_2_FC knockdown measured in HepG2 cells for NPMs (**Methods**). The correlation between accessibility and knockdown was weak but significant (*r* = -0.08, *p* < 10^-17^); however, it became negligible after regressing out DeepRNAi predictions (partial correlation *r* = -0.02, *p* = 0.04). This suggests that DeepRNAi predictions capture much of the accessibility-associated signal in the observed response. Correlations between NPM log_2_FC and GC content or Tm were negligible (|*r*| < 0.01); both attributes correlated with on-target log_2_FC (*n* = 55; GC content: *r* = 0.46, *p* = 3×10⁻⁴; seed Tm: *r* = 0.39, *p* = 0.003). However, regressing out DeepRNAi predictions reduced these to insignificant levels, *p* = 0.49 and *p* = 0.21 respectively, suggesting again that DeepRNAi has learned to capture these signals (**Table S6**).

To investigate whether DeepRNAi learned biophysical attributes *de novo*, that is, without being provided with this information, after the model was finalized we trained a probe on its hidden activations to generate a two-dimensional visualization and predict individual attributes (**Methods**). Although DeepRNAi was not explicitly trained to predict these attributes, its learned representations encode them, as evidenced by smooth variation in attribute values across the two-dimensional projections (**Figure 4b**) and Pearson correlations between probe predictions and measured values on held-out test data. These include normalized expression (*r* = 0.76), eSHAPE accessibility (*r* = 0.38), guide GC content (*r* = 0.93), and seed melting temperature (*r* = 0.91).

The seed region is well known to be critical for siRNA-mRNA hybridization.^10^ To determine whether DeepRNAi has learned this, we assessed the sensitivity of DeepRNAi predictions to individual guide positions by masking each position in turn and measuring the correlation between masked and unmasked predictions. Masking positions 2 through 7 produced a large decrease in correlation, consistent with the known importance of the seed region (**Figure S4**).

We next explored two case studies using published data with biological confounding factors that affected siRNA activity. In the first case study, we compared DeepRNAi predictions to data published by Alnylam for siRNAs targeting human *PLG* (PCT patent application WO2024077148A1, 2024), which included siRNAs spanning both proximal and distal 3′ UTR regions of *NM_000301.5,* the MANE Select (Matched Annotation from NCBI and EMBL-EBI) transcript.^38^

Alnylam’s reported data show consistent reduced on-target activity among siRNAs targeting the distal 3′ UTR downstream of a proximal polyadenylation site, despite complete complementarity to the annotated transcript. DeepRNAi accurately predicted this regional decline in efficacy when supplied with BioFM embeddings, whereas the BioFM-ablated model predicted strong knockdown across the entire 3′ UTR (**Figures 4c,d**). These results demonstrate that BioFM embeddings capture transcript context relevant to alternative polyadenylation and isoform expression, improving the accuracy of DeepRNAi knockdown predictions.

In the second case study, we examined genetic variants that alter RNA processing and siRNA activity. Reference annotations, even if accurate, are often insufficient due to genetic variations that modulate target-site availability in a genotype-dependent manner. Variant filters typically flag only variants overlapping siRNA binding sites, missing non-local regulatory variants that alter mature transcript structure. We hypothesized that BioFM embeddings would allow DeepRNAi to account for these non-local, variant-dependent changes in target-site availability. As a case study, we examined rs10954213 (chr7:128949373:G>A, hg38), a common *IRF5* 3′ UTR variant (global gnomAD^39^ A-allele frequency ∼0.58) that creates a proximal (AAUAAA) polyadenylation signal and increases expression of an isoform lacking approximately 600 bp of the distal 3′ UTR.^40^ *IRF5* is a regulator of inflammatory macrophage and type I interferon signaling, and siRNA knockdown reduced inflammatory signaling and improved infarct healing in a murine model.^41–43^

We first confirmed that the BioFM Enigma^44^, which is used by DeepRNAi, predicted RNA-seq signal across the shared transcript region and an abrupt loss of signal downstream of the created polyadenylation site when provided with the variant, which is consistent with increased expression of the shortened isoform. Consequently, when scored in the rs10954213 A-allele context, DeepRNAi predicted weaker knockdown for all siRNAs targeting the missing ∼600 bp segment of the distal 3′ UTR (**Figure 4e**). In contrast, the BioFM-ablated model only predicted differences in activity for siRNAs directly overlapping the variant, as it can only account for local context. A variant-aware design pipeline should therefore look beyond disruptions in the target binding site to select molecules that retain activity across common genotypes.

These results show that through BioFMs, DeepRNAi has learned at least part of the biology of siRNA activity and can capture activity effects arising from isoform expression differences and regulatory variants. This enables accurate virtual screening of candidate siRNAs across complex genetic contexts, including non-local variant effects on transcript structure, and allows predictions to be individualized for specific populations.DeepRNAi prioritizes Inclisiran in a retrospective PCSK9 design analysis

Having established that DeepRNAi accurately predicts siRNA effects across unseen guides, targets, cellular contexts and species, we next asked whether it could identify potent and specific sequences selected for preliminary in vitro screening.

We first evaluated DeepRNAi in a conventional tiling and selection workflow using human *PCSK9* as a target. To prevent direct target leakage, DeepRNAi was retrained after excluding all siRNAs targeting *PCSK9*. We then tiled 3,615 fully complementary siRNAs across the most highly expressed *PCSK9* transcript in liver (NM_174936.4) and jointly prioritized them according to predicted on-target activity and aggregate predicted off-target burden (**Methods**). Filtering for greater than or equal to 75% predicted knockdown reduced the pool to 10 candidates, of which six also had low predicted off-target burden (≤ 0.08, as calibrated from **Figure 3e**). The FDA approved therapeutic Inclisiran was among this final set of candidates. DeepRNAi predicted approximately 76% on-target knockdown for Inclisiran together with low off-target burden and only one NPM transcript predicted to be differentially expressed (**Figure 5a**). Across the complete candidate pool, predicted potency was correlated with increasing off-target burden (*r* = -0.295), demonstrating that the strongest predicted molecules were also generally less selective. This target-held-out analysis shows that DeepRNAi prioritized screening of a highly specific clinically successful siRNA.

**Figure 5:**
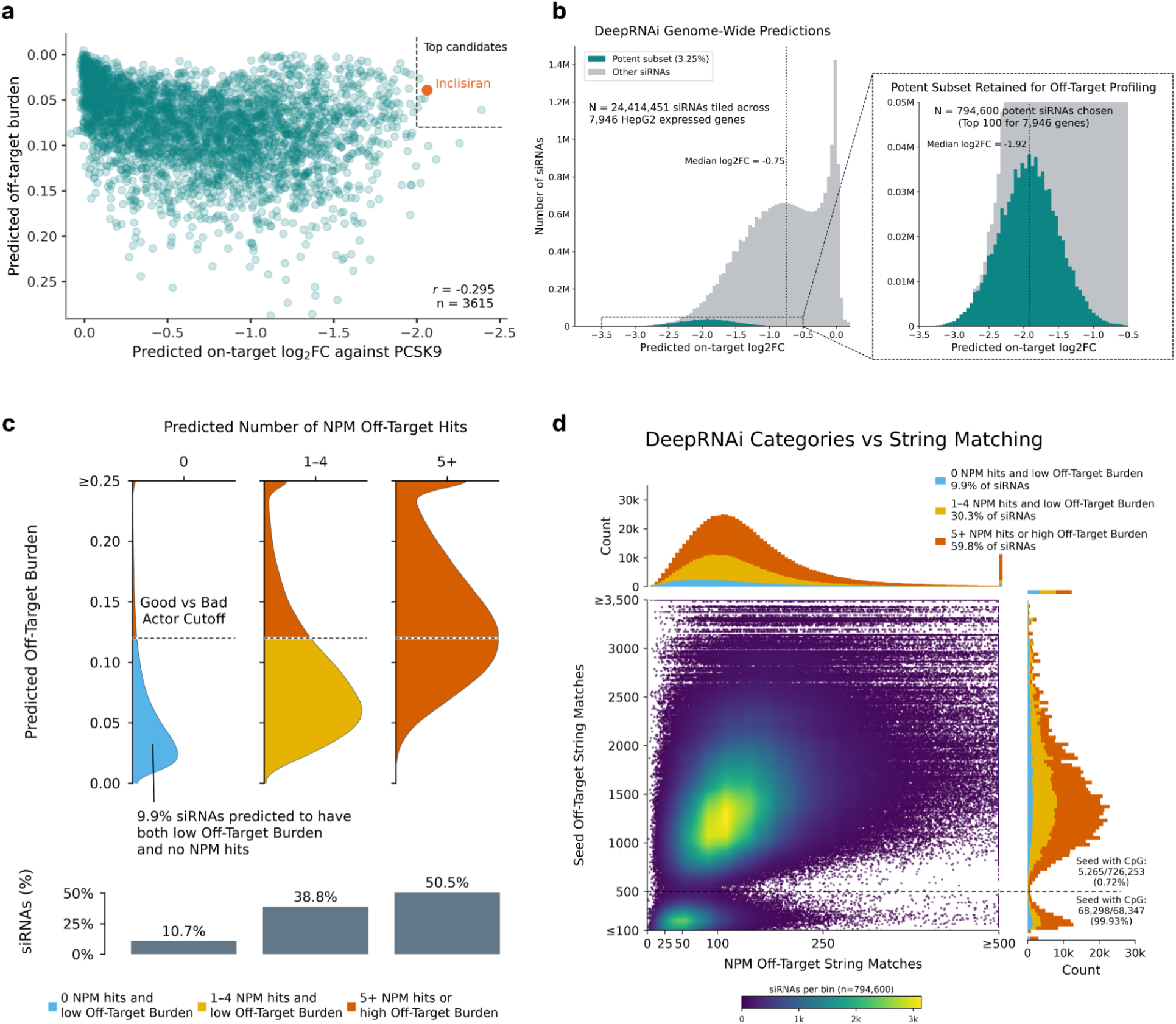
DeepRNAi predictions for PCSK9 and the full genome. **a)** Predicted on-target log_2_FC and off-target burden for all siRNAs tiling PCSK9. Off-target burden is the difference in mean log_2_FC (restricted to genes with log_2_FC < 0) between transcripts without an NPM or seed match and those with. The approved therapeutic Inclisiran shown with an orange dot. Top candidates are siRNAs with predicted on-target log_2_FC ≤ -2 and predicted off-target burden ≤ 0.08. The axes have been inverted such that stronger on-target log_2_FC is toward the right, and higher off-target burden is toward the bottom. **b)** Distribution of predicted on-target knockdown across all genome-wide siRNAs, the 100 most potent siRNAs per gene are shown in cyan (3.25%). **c)** Off-target profiles among the 794,600 prioritized siRNAs: 9.9% had no active NPM-matched off-targets and low predicted burden (< 0.12; threshold from the rodent hepatotoxic/FDA approved siRNA dataset shown in **Figure S3c**), 30.3% had 1–4 active NPM-matched off-targets and low burden, and 59.8% had ≥5 active NPM off-targets or high burden. **d)** NPM and seed matches per siRNA across the 794,600 potent siRNAs, with marginal histograms by off-target category. A distinct cluster with fewer than 500 seed matches almost entirely contained siRNAs with CpG dinucleotides in the guide seed region (99.93% vs 0.72% among siRNAs with ≥500 seed matches, ∼140-fold enrichment).

### A genome-wide map of siRNA off-target burden

Given the ability to retrospectively prioritize Inclisiran, we extended a similar analysis genome-wide to assess the overall space of potent molecules with favorable off-target specificity. We first tiled all 7,946 protein-coding transcripts we found to be sufficiently expressed in HepG2 and scored 24.4 million candidate siRNAs for on-target potency.

To conduct off-target burden analysis, we retained the top 100 most potent siRNAs for each gene (3.25%), to simulate molecules that might advance to experimental screening. These had a median predicted on-target log_2_FC of −1.92 (**Figure 5b**). We then used DeepRNAi to predict the knockdown of off-target transcripts containing NPMs or seed matches. Only 78,574 siRNAs (9.9%) were predicted to have both zero active off-target NPMs and low transcriptome-wide off-target burden, constituting a “DeepRNAi-clean” profile (**Figure 5c**) (**Methods**). Across the genes in this set, we found an average of ∼10 DeepRNAi-clean siRNAs amongst the top 100 per gene (standard deviation of 4.5). The DeepRNAi-clean siRNAs were not well-separated by string matching methods (**Figure 5d**). For example, using a strict filter that requires less than 5 NPM sequence matches and less than 500 seed matches recovered only 13 of these 78,574 predicted clean siRNAs (0.02%), representing only 13 of the 7,946 target genes. This filter also retained 37 siRNAs with high predicted off-target burden. These findings suggest that jointly modeling potency and off-target activity to prioritize clean and potent siRNAs for experimental evaluation could reduce costly late-stage failures or program resets, preempt potential regulatory delays due to NPM and seed match hits, and vastly reduce the amount of screening needed to find good drug candidates.

Predictions using DeepRNAi for the 100 most potent, anonymized siRNAs for any gene expressed in HepG2 cells along with full results and downloadable data for PCSK9 are available through RNAi Explorer (https://tools.deepgenomics.com/rnaiexplorer/).

## Discussion

By acting upstream of protein production, siRNA therapeutics can address targets that other modalities cannot, including proteins lacking a druggable binding pocket and intracellular targets inaccessible to antibodies. This can make them safer, more potent and longer lasting than many drugs currently on the market. Since they are designed from a discrete set of sequences and chemical modifications, they are “programmable” and uniquely suited to discovery using computational modeling and AI. However, each siRNA can potentially bind to a large number of off-target mRNA sequences with only a small proportion actually affected, so current discovery workflows proceed with an inefficient step-wise process of selection by on-target potency followed by screening for genome-wide impacts with RNA-seq and animal studies. This can result in the best candidates being dropped early while candidates with predictable liabilities advance to animal models. Current bioinformatics and machine learning approaches cannot accurately predict these effects, and until now there has not existed a comprehensive public benchmark dataset to evaluate predictive methods.

This work establishes a benchmark dataset and a model showing for the first time that transcriptome-wide responses to siRNAs can be predicted across unseen siRNA sequences, target genes, screening conditions, cell lines, genotypes and species, with performance that approaches the reproducibility of independent experiments measured by RNA-seq. The performance of DeepRNAi suggests it has learned transferable rules of hybridization-dependent RNAi repression. *In silico* screening can be used to prioritize higher-quality candidates early on and mitigate costly redesign cycles and late-stage failures driven by liabilities discovered only after lead selection. The model captures intended target knockdown, near-perfect-match (NPM) effects and microRNA-like seed-match effects, and outperforms TargetScan and traditional machine learning baselines by a wide margin. Because DeepRNAi uses BioFM embeddings to capture relevant genome context, it can adjust for inaccurate transcript annotations, accurately predict off-target effects for genomic variants, and adapt to new species or patient contexts.

This points the way to a more streamlined discovery process where all possible siRNA guides (including potentially those with nucleotide substitutions and/or chemical modifications) are assessed *in silico* for both on-target potency and off-target liabilities across many cellular contexts, genetic backgrounds and species. This would greatly increase the number of high quality candidate molecules, widening the lead-selection funnel, and increasing the probability that those leads will progress through later stages of development. This approach would also allow siRNA drug discovery programs to potentially replace slow and error-prone experimental cycles with fast, computational cycles, where benefits and risks at each step along the path to the clinic can be predicted and controlled. Such a preclinical pipeline could allow for systematic discovery of therapies for a larger set of diseases including those with smaller population sizes that are currently underserved.

Because off-target prediction remains understudied relative to on-target potency prediction, the field lacks a standardized public benchmark. Existing transcriptome-wide siRNA datasets are scattered across individual publications, use heterogeneous chemistries, cell systems, dosing and controls, and have not been assembled into a common evaluation set. Therefore, to help support model development and facilitate consistent evaluation, we are releasing a public dataset of 240 siRNAs, partitioned into 164 for training and 76 for testing (**Supplementary Materials**, model performance on this set shown in **Figure S5** and **Table S7**). We invite the community to use this resource to accelerate the development of future models for siRNA on- and off-target prediction.

Going forward, we envision several ways to improve DeepRNAi. First, as we continue to screen different siRNAs and add them to our training set, our scaling curves (**Figure 2c**) suggest that further predictive performance gains are achievable. Second, recent work by several groups has highlighted the importance of targeted chemical modifications to improve on-target efficacy while reducing off-target burden.^45–47^ Adding siRNAs with diverse chemical modifications to model training should not only allow future versions of DeepRNAi to better predict efficacy across a wider range of siRNA chemistries, but to generalize to novel modifications, further increasing the breadth of the design space to ultimately find better molecules. Third, incorporating training data and benchmarking metrics from *in vivo* screens and clinical trial outcomes will further augment the utility of DeepRNAi in drug discovery pipelines. Since the ultimate goal is to design siRNA therapies for the clinic, a prospective evaluation of DeepRNAi’s performance for siRNA design on candidates that are advanced through a drug-discovery pipeline will be the clearest test.

In summary, DeepRNAi is a highly effective tool for siRNA design, potentially streamlining *in vitro* screening, and allowing drug developers to identify a greater number of potent siRNAs with limited off-target burden earlier in the screening process. By rapidly prioritizing higher quality molecules up front and decreasing the risk of toxicity signals and regulatory complications, we believe this model is capable of accelerating the development of the next generation of siRNA therapeutics.

## Methods

### Definitions and parameters

#### Near-perfect match (NPM)

An siRNA-mRNA pair with one or more sites of high complementarity (edit distance < 2) between the mRNA and positions 2 through 16 (one indexed) on the siRNA guide strand.

#### Seed match

An siRNA-mRNA pair in which the mRNA’s 3′ UTR contains at least one site with perfect complementarity to positions 2-7 of the siRNA guide strand. Seed-match sites are classified into four types:

- **6mer:** Complementarity to guide positions 2–7.
- **7mer-A1:** Complementarity to guide positions 2–7, with an adenosine in the mRNA opposite guide position 1.
- **7mer-m8:** Complementarity to guide positions 2–8.
- **8mer:** Complementarity to guide positions 2–8, with an adenosine in the mRNA opposite guide position 1.

Sites satisfying multiple definitions are assigned the highest-ranking type: 8mer > 7mer-m8 > 7mer-A1 > 6mer.

#### Hit

An siRNA-mRNA pair with log_2_FC < -0.5, or ∼30% knockdown (unless otherwise specified).

### Data processing for evaluations

HepG2 (40k per well), Huh7 (30k per well), and Hep3B (30k per well) cells were transfected with either siRNA or Mock samples using RNAiMAX reagent at 0.3 μL/well. All siRNA samples were transfected at a 10 nM dose in triplicate. After 48 hours, cells were lysed and purified RNA was sent for RNA Sequencing.

RNA-Seq was performed using the Illumina stranded mRNA sequencing approach (Illumina, San Diego, CA; cat. no. 20040534) and short-read Illumina 3′ end sequencing (Plasmidsaurus, https://plasmidsaurus.com/technical-documentation/rna). For 3′ end sequencing the libraries were sequenced on the Illumina NovaSeq X Plus platform using asymmetric paired-end sequencing, which generates roughly 10 million deduplicated 3′ end reads. The mRNA reads were approximately 90 bases in length and enriched near the 3′ end of captured transcripts. Stranded mRNA sequencing was performed on the Illumina NovaSeq X Plus instrument with up to 25B reads per flow cell and ∼20M-25M reads per sample. Reads were trimmed for quality and adapter content using fastp^48^ (v0.23.2). Sequencing quality was assessed using MultiQC^49^ (v1.22.2) with FastP and FastQC^50^ (v0.11.9) outputs used as input. Trimmed reads were pseudo-aligned to the reference transcriptome (GRCh38.p14 Refseq v110) with Kallisto^51^ (v0.46.2) using 100 bootstrap samples for transcript abundance estimation.

Kallisto-derived abundance estimates were processed in R^52^ (v4.6.1) and aggregated to the gene level. Within each experiment, genes with a median TPM > 1 across Mock samples were retained, and samples with fewer than 5 million reads were excluded. Differential expression analysis was performed using DESeq2^53^ (v1.52.0), fitting a separate model for each siRNA within each sequencing experiment using the corresponding Mock samples as negative controls. Condition (case vs. control) was included as the explanatory variable, with sequencing run included as an additional covariate when replicates spanned multiple runs. Differential and stable expression was assessed using separate hypothesis tests (altHypothesis = "greaterAbs", lfcThreshold = 0; and altHypothesis = "lessAbs", lfcThreshold = 0.4, respectively). P-values were adjusted using the Benjamini-Hochberg procedure,^54^ and log₂ fold-change estimates were shrunk using the adaptive shrinkage method implemented in ashr.^55^

PacBio long-read sequencing data, initially provided in unaligned BAM format, were converted to FASTQ format with bam2fastq (v3.5.0) from the PacBio BAM toolkit (pbtk)^56^ and concatenated across SMRT Cells. The resulting reads were aligned to the GRCh38.p14 reference genome using minimap2^57,58^ (v2.30-r1287) in splice-aware mode with secondary alignments suppressed discarding multi-mapped reads (-ax splice -uf --secondary=no --MD).

All DeepRNAi predictions are based on training on pre-shrinkage log_2_FC values from DESeq2.^53^ As such, all evaluations are also based on these pre-shrinkage log_2_FC values. For our held-out test set, to ensure that models were not simply memorizing the mean response of each transcript, evaluations were performed on mean-centered pre-shrinkage log_2_FC values. Specifically, for each transcript and cell type combination, we calculated the mean pre-shrinkage log_2_FC using the train set, and subtracted this value from the corresponding pre-shrinkage log_2_FC values in the test set. The resulting log_2_FC value from this subtraction is the target in all of our evaluations performed on our held-out test set. For all other RNA-seq datasets, no subtraction of the mean effect was performed and the pre-shrinkage log_2_FC values were used as the evaluation target.

Within each cell line for each dataset, a single transcript isoform corresponding to the highest expressed transcript for each gene was selected to represent the gene, as determined by the mean TPM values of Mock samples. This mean TPM value is referred to as the baseline expression of the transcript and was used for further filtering (see below). All annotations pertaining to NPMs and seed matches were made with respect to the selected transcript isoform. Only protein-coding transcripts with a baseline expression of at least 10 TPM were included in the evaluations, with the exception of transcripts for genes targeted by siRNAs in our dataset, which were retained if they had a baseline expression of at least 1 TPM. Note that these filters resulted in the exclusion of 2 genes (INHBE and SULT1A4) from on-target evaluations within Huh7 cells.

Transcriptome-wide differential expression data generated from RNA-seq performed on siRNA-treated samples will characterize both *direct* events, consisting of siRNA-driven interactions of RISC with an mRNA transcript, resulting in downregulation of that transcript’s expression, and *indirect* events, encompassing upregulation or downregulation events in an siRNA-treated sample relative to a control that do not arise as an immediate consequence of RISC-transcript interactions. These indirect events may stem from a multitude of sources, including stress response to siRNA treatment and regulatory cascades through the gene regulatory network (GRN). Some indirect events are downstream consequences of direct interactions mediated through the GRN, while others are entirely independent of direct events. Before training DeepRNAi, we applied an unsupervised, sequence-agnostic preprocessing procedure to reduce shared response structure across perturbations. The procedure did not use siRNA sequence, siRNA-mRNA complementarity or binding-site annotations. The resulting values were used as training targets, while model evaluation was performed against experimentally measured log2 fold changes.

### DeepRNAi model training

DeepRNAi was implemented in PyTorch and trained for eight epochs using the Huber loss (delta = 4) and the AdamW optimizer (weight decay = 0.05). The batch size was 512, and gradient clipping with a maximum norm of 1 was applied to stabilize training. The learning rate followed a linear warm-up schedule, up to a maximum of 0.0001, followed by cosine annealing. We trained on the mean-centered (defined above) pre-shrinkage log_2_FC values. Details of the model architecture are proprietary and are not disclosed.

### Random forest training

The random forest regression model from Scikit-learn^59^ was trained on the same internal Deep Genomics data as DeepRNAi. First, the training data were filtered for siRNA-transcript pairs that have a seed or NPM binding site.

A single binding site per transcript was selected based on the eCDF distribution of log_2_FC values, according to the following rank order: NPM with an edit distance of 0, 1, or 2; 8mer seed; NPM with an edit distance of 3; 7mer-m8 seed; 7mer-A1 seed; 6mer seed.

A total of 21 features were extracted as model input: 8 binary features encoding binding site type, categorical features for chemistry and cell type, 2 siRNA guide sequence-based continuous features (19mer GC content and 7mer seed Tm), 3 binding site continuous features (GC content, PhastCons conservation scores, and guide siRNA edit distance), and 6 transcript-level continuous features (mean TPM of control samples, 3′ UTR length, ORF length, transcript GC content, near-perfect complementarity match count, and seed match count).

### String matching model

The string matching approach uses the same internal training data as DeepRNAi to obtain the mean experimental log_2_FC for each NPM edit distance and seed type. First, the training data is filtered for off-target protein coding transcripts with a baseline TPM expression greater than or equal to 10. This model is susceptible to many ties and ranks siRNA according to the following rank order: NPM with an edit distance of 0, NPM with an edit distance of 1, 8mer seed, NPM with an edit distance of 2, 7mer-m8 seed, 7mer-A1 seed, and 6mer seed. The background set consists of all other transcripts without an NPM or seed hit.

### Seed melting temperature model

A baseline model for 7mer seed Tm is used to benchmark performance of off-target transcripts that have a seed hit. The Tm of the siRNA guide strand positions 2 to 8 is acquired using the Primer3 package.^60^

### Statistical analysis

Bootstrap analysis was used to quantify significance between on-target Pearson correlation values, and NPM and seed average precision. To do so, we randomly sampled siRNAs with replacement, and calculated the associated metrics. We repeated this procedure 2,000 times, using the same resampled siRNAs for all methods within each comparison. To calculate an approximate two-sided p-value for the null hypothesis of no difference between two values, we calculated the proportions of paired bootstrap differences that were nonpositive and nonnegative, then doubled the smaller proportion. P-values were capped at 1.

### Average precision

We used average precision (AP) to evaluate the identification of functional knockdown among siRNA-mRNA pairs with NPM or seed matches. Pairs with observed, mean-subtracted log_2_FC < −0.5 and adjusted p-value < 0.05 were labelled positive. Pairs with log_2_FC < −0.5 and adjusted p-value > 0.05 were excluded; all remaining pairs were labelled negative. We calculated AP using scikit-learn’s *average_precision_score*, with the negative of predicted log_2_FC values as scores so that stronger predicted knockdown received a higher score.

### Decoding model attributes

To assess whether biophysical attributes (normalized expression, eShape accessibility, guide GC content and seed Tm) could be decoded from DeepRNAi’s internal representations, we trained a decoder on activations from a late-stage DeepRNAi layer. These activations passed through an MLP with two hidden layers of 32 units each and GELU activations, followed by a projection to a two-dimensional bottleneck. Activity from the bottleneck layer passed through a second MLP with two hidden layers of 32 units each and GELU activations, and finally a linear layer mapped these values onto attribute predictions. A stop-gradient operation immediately after the DeepRNAi activations prevented decoder training from updating DeepRNAi weights. Decoders were trained and evaluated using the same training and held-out test splits used throughout the study.

### Reproducibility of RNA-seq measurements

We assessed cross-cell-line RNA-seq reproducibility using 11 held-out siRNAs screened independently in HepG2 and Huh7 cells. We matched siRNA-mRNA pairs across cell lines and used observed responses in one cell line to define functional knockdown labels and observed responses in the other as prediction scores. In the reference cell line, siRNA-mRNA pairs were classified as functional as defined above.

When comparing model accuracy to the reproducibility of RNA-seq measurements, we compared their baseline-subtracted APs. The motivation is that if method A achieves an AP of 0.5, and method B achieves an AP of 0.4, and the baseline is 0.3, then it would be misleading to state that method B achieves 80% (0.4 / 0.5) of the accuracy of method A. Thus, we subtracted the baseline from both values before taking the ratio. Thus, in our example above, we state that method B achieves 50% of method A ((0.4 - 0.3) / (0.5 - 0.3)).

To assess reproducibility within cell lines (**Table S4**), we wanted to maximize statistical power. Thus, we trained a new version of DeepRNAi in which we moved all 14 siRNAs screened multiple times in the same cell line from the training to the test set, and re-evaluated. Since there already existed a single siRNA screened multiple times in the standard test set, this results in a held-out set of 15 siRNAs.

### Off-target burden

For DeepRNAi predictions, “off-target burden” for a given siRNA was calculated as the difference in predicted mean log_2_FC, after filtering to just predictions that were less than zero log_2_FC, between the set of transcripts that do not contain an NPM or seed match to that siRNA and the set of transcripts with an NPM or seed matches to that siRNA. The metric as calculated from experimental RNA-seq data was calculated in the same way, but using the mean experimental pre-shrinkage log_2_FC values.

For the genome-wide designs (**Figure 5**), because we only computed predictions for off-target transcripts with a seed match or NPM match, we used a modified metric for off-target burden. This metric just took the mean of the predicted log_2_FC values that were less than zero for transcripts that contained an NPM or seed match. We used a cutoff of 0.12 for this value based on the rodent hepatotoxic/FDA approved siRNA dataset in **Figure S3c**.

### Transcript selection for the rat toxicity evaluation

For DeepRNAi predictions, target transcripts were selected as the most highly expressed transcript for a given gene in that cell line (details on how this transcript is selected is described in the **Data processing for evaluations** section above). This ensures that the set of transcripts used to compute off-target burden are concordant between both the experimental and *in silico* data.

### Evaluation against rat *in vitro* and *in vivo* data

Publicly available rat liver and primary rat hepatocyte RNA-seq data, in the form of gene-level raw count matrices and precomputed differential gene expression results from Schlegel et al.^17^ was downloaded from the NCBI Gene Expression Omnibus^61^ (GEO; accession GSE184929). The log_2_FC values correspond to the DESeq2 beta-prior-shrunken estimates^53^. For each gene, the most principal transcript (RNor_6.0, Ensembl release 88) as per APPRIS,^62^ was selected as the representative transcript for the gene. Genes without an APPRIS transcript annotation were excluded. Guide and passenger strand sequences for siRNAs were obtained from Supplementary Table S1 of Schlegel et al. and matched to the RNA-seq data using siRNA identifiers.

To restrict analyses to genes with sufficient baseline expression, approximate TPM values were calculated from the raw gene-level counts of control samples. Gene counts were normalized by the total exonic length of the corresponding transcript and scaled to one million RPKM units within each sample. These TPM values were averaged across biological replicates within each control group, and the median control TPM across rat experimental groups was calculated for each transcript. Only transcripts within the top 33% of median baseline expression were retained. Genes lacking matched control expression measurements were excluded.

Evaluations were done including two siRNAs from the study: D1 and D6. For evaluations against the *in vivo* rat liver data, we show the result of a “predictor” that combines the *in vitro* primary rat hepatocyte differential gene expression results with DeepRNAi predictions. This predictor takes the absolute value of DeepRNAi predictions and multiplies it with the log_2_FC value of the same siRNA and readout transcript pair from the *in vitro* primary rat hepatocyte experiment. Multiple methods of combining (e.g., averaging, min, and max) were explored, but we found that this was the best performing method.

## Acknowledgements

The authors thank M. Edbrooke, A. Gao, A. Gouveia, C. Hart, Y. LeCun, D. Lim, L. MacPhie, J. Maraganore, G. McAllister, R. Meyers, V. Myer, B. Neale, N. Pursell, U. Saxena and R. Scheller for conversations about the science, the manuscript and the publication strategy.

## Author contributions

N.Y.M., V.S.C., H.Z., B.K., and J.L. designed the model architecture and implemented DeepRNAi. T.T.Y.L., J.W., M.S. and R.L. developed the benchmark datasets and evaluation framework. N.Y.M., T.T.Y.L., J.W., R.L. and A.C. performed computational analyses and generated figures. B.F., K.A., S.J. and W.T.L. generated and analysed the RNA-seq data. A.C., N.H.L., S.D.F. and B.J.F wrote the manuscript. N.L.C., E.B. and C.B.C. provided scientific and analytical input. A.C., N.H.L., S.D.F., G.R.H. and B.J.F. conceptualized and supervised the work. All authors reviewed and edited the manuscript.

## Supplementary Information

### Data availability

In order to establish the machine learning task of predicting off-target effects for siRNA treatments we present here a dataset that researchers can use to benchmark methods and compare directly to the work presented above. This consists of 1.84M siRNA-mRNA pairs from 240 siRNAs. For benchmarking purposes, we split that data into a public train set of 164 siRNA targeting the genes *TRUB1*, *EIF1AX*, *NCL*, *INHBE* and *SULT1A4* and a public test set of 76 siRNA targeting the genes *ZHX2*, *SLC29A1*, *EDEM2*, and *APOC1*. These train and test sets are roughly balanced across cell lines (HepG2 and Huh7), siRNA format and chemistry and can be additionally split for cross-cell line prediction tasks. The data is hosted at https://github.com/deepgenomics/DeepRNAi.

### Mini-DeepRNAi model performance

We trained a version of DeepRNAi, termed “mini-DeepRNAi,” exclusively on the public training set of 164 siRNAs. We evaluated its performance on the public test set consisting of 76 siRNAs alongside a Random Forest model and a version of mini-DeepRNAi without BioFM embeddings, both trained on the same public training set (**Figure S5**, **Table S7**)The mini-DeepRNAi significantly outperformed the RF and the mini-DeepRNAi model without BioFM embeddings across NPMs and seed matches, whereas the

### Supplementary Figures

**Figure S1:**
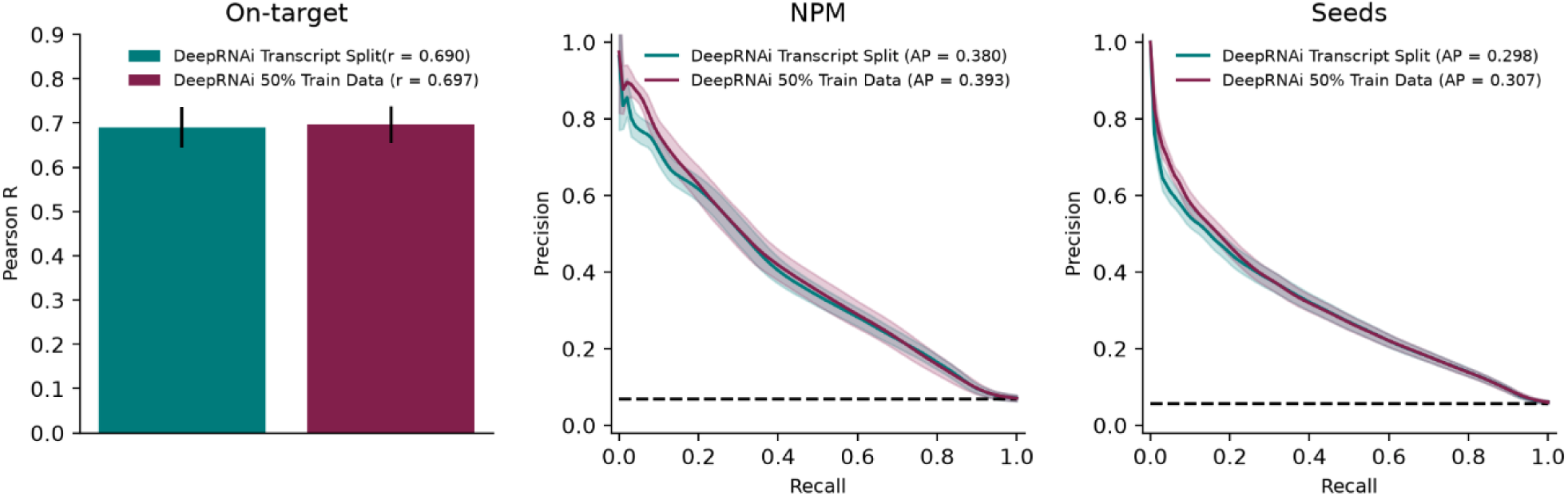
DeepRNAi performance on 50% data. Comparing DeepRNAi model trained on half the readout transcript measurements (as shown in Figure 2b) versus a DeepRNAi model trained on 50% of siRNA-mRNA pairs in order to match the quantity of training data. Performance metrics are the same as those shown in Figure 2.

**Figure S2:**
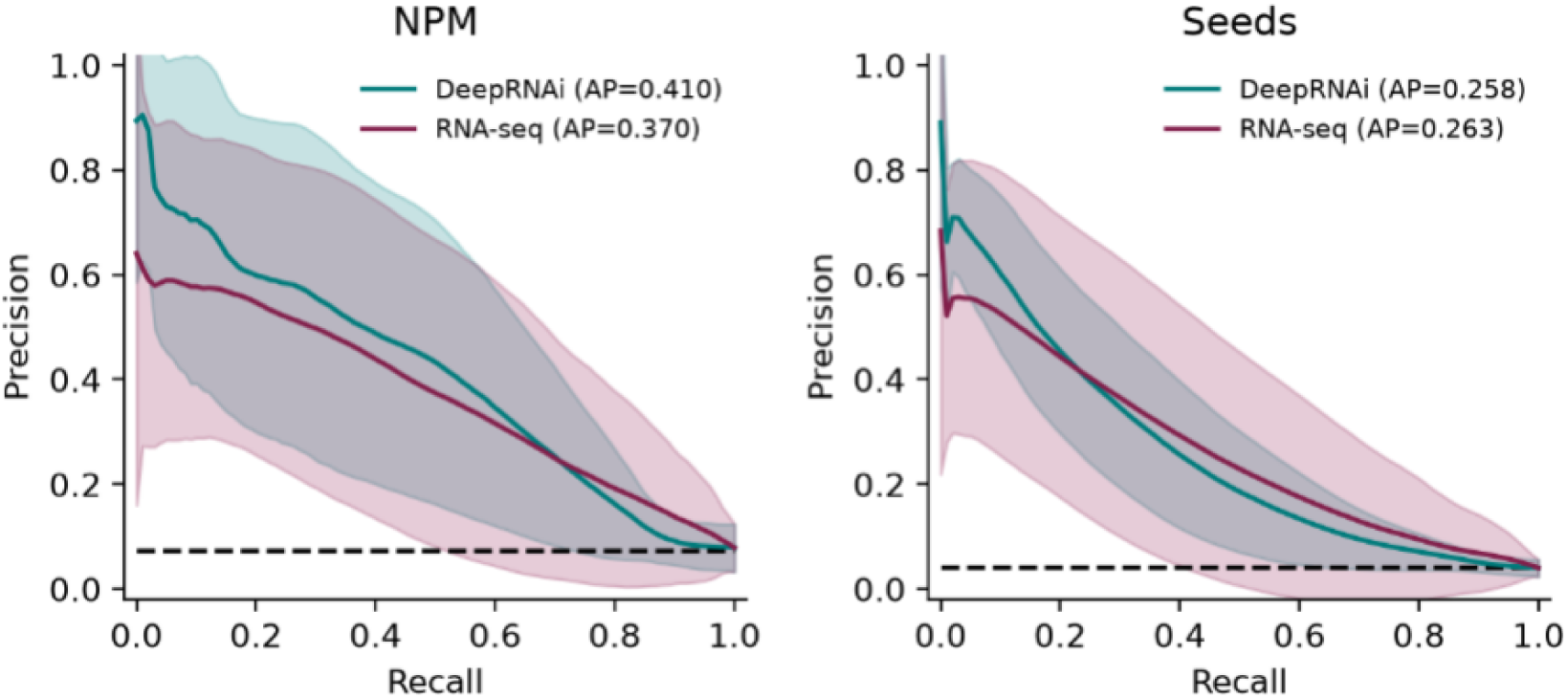
DeepRNAi accuracy compared to RNA-seq within cell-line reproducibility. Comparing DeepRNAi predictions against RNAseq reproducibility for 15 siRNA that were screened twice within the same cell line. Performance metrics are the same as those shown in Figure 3a.

**Figure S3:**
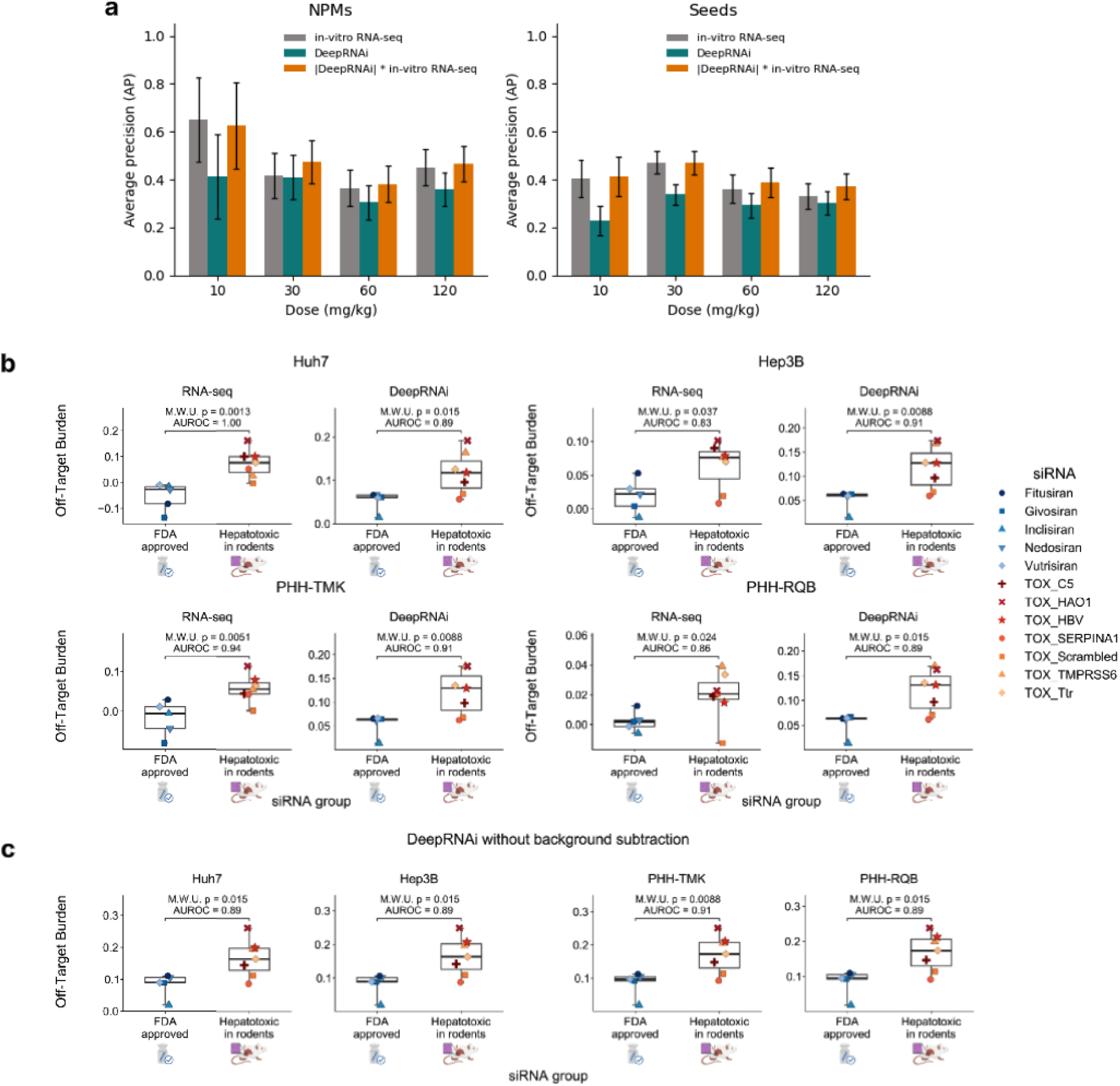
DeepRNAi performance on *in vitro* and *in vivo* test data. **a)** Bar plots showing the average precision (AP) of DeepRNAi, *in-vitro* RNA-seq done in rat hepatocytes, and a combination of the two for classifying hits (log_2_FC < -0.5 with adjusted p-value < 0.05) from non hits (log_2_FC > -0.5) amongst NPMs (left) and seed matches (right) for four different doses of two siRNAs from Schlegel et al. 2022.^17^ The error bars represent ± one standard deviation of the AP calculated across 10,000 bootstraps. **b)** Box plots showing off-target burden between approved therapeutic siRNA and siRNA believed to show rat toxicity in Janas et al.^7^ Off-target burden defined as the difference in mean log_2_FC (for log_2_FC < 0) between transcripts without NPM or seed matches and transcripts with matches. **c)** Box plots showing predicted off-target burden from DeepRNAi for the same siRNAs and cell lines in b), but with a metric that is defined as just the mean predicted log_2_FC (for log_2_FC < 0) multiplied by -1.

**Figure S4:**
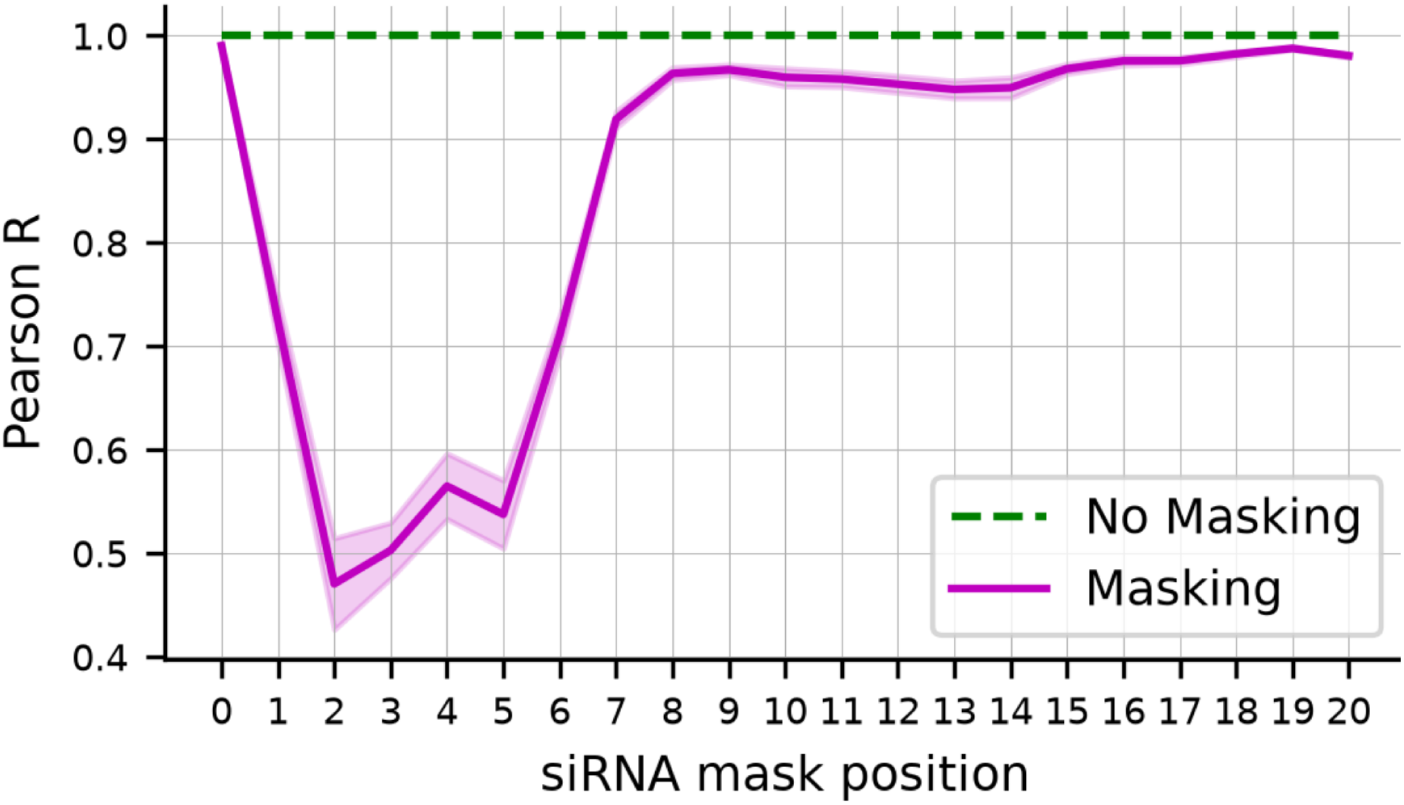
Sensitivity of DeepRNAi to individual guide positions. The Pearson correlation between DeepRNAi predictions with a single guide position masked and the unmasked predictions are shown for each position. Error bars show +/- one standard deviation. Note that positions are zero-indexed.

**Figure S5:**
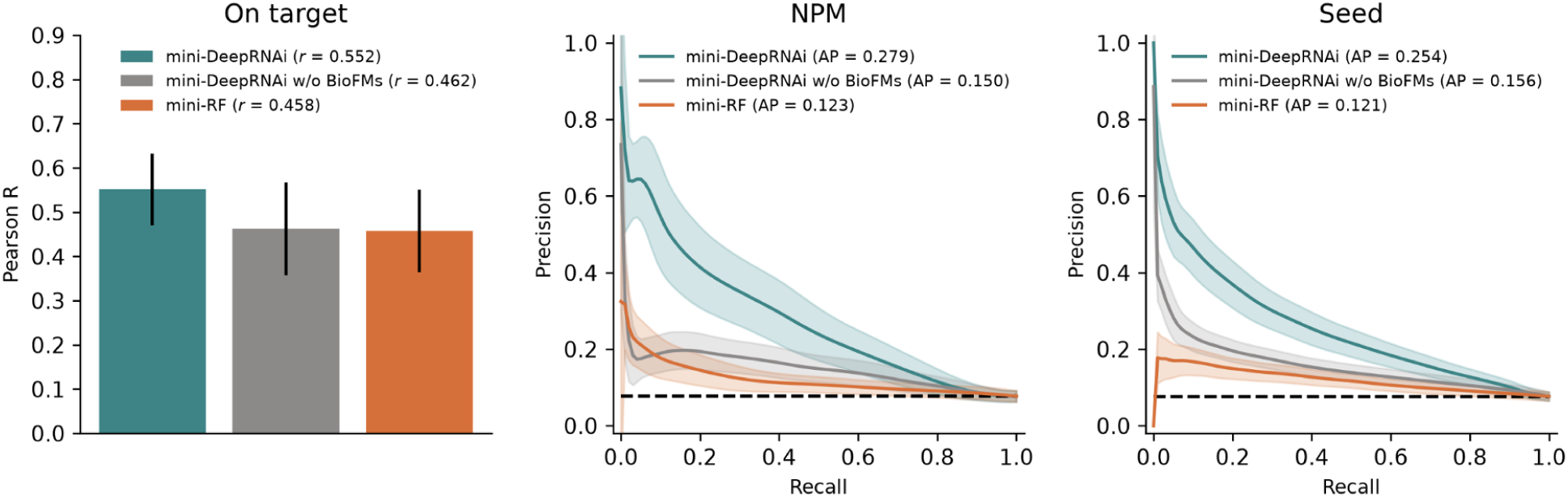
mini-DeepRNAi performance. Performance of models trained and evaluated on the public data set (164 training, 76 test siRNA). Pearson correlation for on-target knockdown and precision-recall curves (PRCs) with average precision (AP) for classifying NPM and seed match knockdown (log_2_FC < -0.5 and adjusted *p* < 0.05) for test siRNA targeting genes that were not seen in the training data. Metrics are shown for DeepRNAi, a version of DeepRNAi that was trained without BioFMs (DeepRNAi w/o BioFMs) and a random forest model (RF). The error bars represent ± standard deviation.

### Supplementary Tables

**Table S1:** DeepRNAi and competitor performance on predicting held-out siRNAs.

|  | **On-target** |  | **NPMs** |  | **Seed matches** |  |
|---|---|---|---|---|---|---|
| **Model** | Pearson correlation | p-value vs DeepRNAi | Average Precision | p-value vs DeepRNAi | Average Precision | p-value vs DeepRNAi |
| DeepRNAi | **0.736** | - | **0.424** | - | **0.337** | - |
| DeepRNAi w/o BioFMs | 0.517 | 0.004 | 0.297 | <0.001 | 0.191 | <0.001 |
| Random Forest | 0.570 | 0.021 | 0.215 | <0.001 | 0.176 | <0.001 |
| String matching | - | - | 0.106 | <0.001 | - | - |
| TargetScan | - | - | - | - | 0.079 | <0.001 |
| SeedTM | - | - | - | - | 0.047 | <0.001 |
| Baseline rate | - | - | 0.070 | - | 0.060 | - |

**Table S2:** DeepRNAi and competitor performance on predicting held-out siRNAs and transcripts.

|  | **On-target** |  | **NPMs** |  | **Seed matches** |  |
|---|---|---|---|---|---|---|
| **Model** | Pearson correlation | p-value vs DeepRNAi | Average Precision | p-value vs DeepRNAi | Average Precision | p-value vs DeepRNAi |
| DeepRNAi | **0.691** | - | **0.379** | - | **0.298** | - |
| DeepRNAi w/o BioFMs | 0.554 | 0.043 | 0.300 | 0.026 | 0.203 | <0.001 |
| Random Forest | 0.543 | 0.033 | 0.195 | 0.002 | 0.119 | <0.001 |
| Baseline rate | - | - | 0.070 | - | 0.060 | - |

**Table S3:**
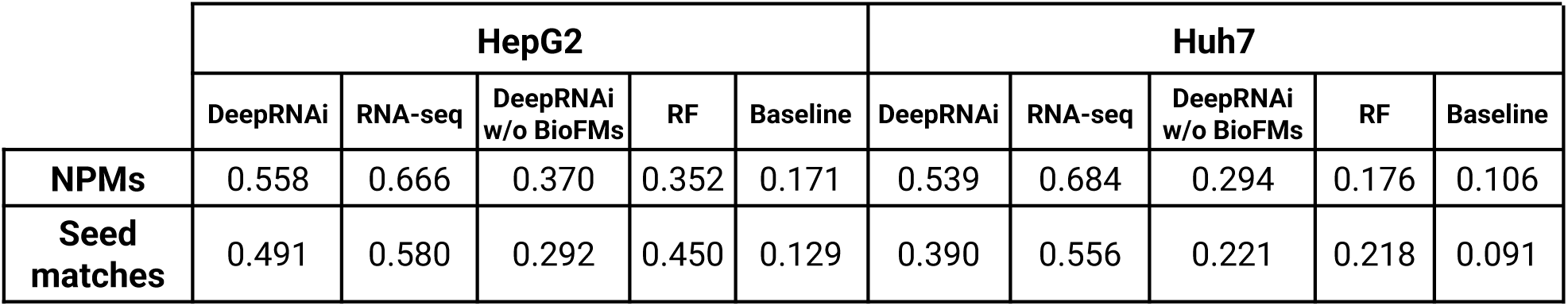
DeepRNAi accuracy compared to RNA-seq cross-cell-line reproducibility. Average precision of DeepRNAi versus observed cross-cell-line RNAseq measurements for NPMs and seed matches on 11 siRNAs screened in both HepG2 and Huh7.

|  | HepG2 |  |  |  |  | Huh7 |  |  |  |  |
|---|---|---|---|---|---|---|---|---|---|---|
|  | DeepRNAi | RNA-seq | DeepRNAi w/o BioFMs | RF | Baseline | DeepRNAi | RNA-seq | DeepRNAi w/o BioFMs | RF | Baseline |
| **NPMs** | 0.558 | 0.666 | 0.370 | 0.352 | 0.171 | 0.539 | 0.684 | 0.294 | 0.176 | 0.106 |
| **Seed matches** | 0.491 | 0.580 | 0.292 | 0.450 | 0.129 | 0.390 | 0.556 | 0.221 | 0.218 | 0.091 |

**Table S4:** DeepRNAi accuracy compared to RNA-seq within cell-line reproducibility. Average precision of DeepRNAi versus observed within-cell-line RNA-seq measurements for NPMs and seed matches on 15 siRNAs screened twice within a single cell line.

|  | **DeepRNAi** | **RNA-seq** | **Baseline** |
|---|---|---|---|
| **NPMs** | 0.410 | 0.370 | 0.073 |
| **Seed matches** | 0.258 | 0.263 | 0.040 |

**Table S5:**
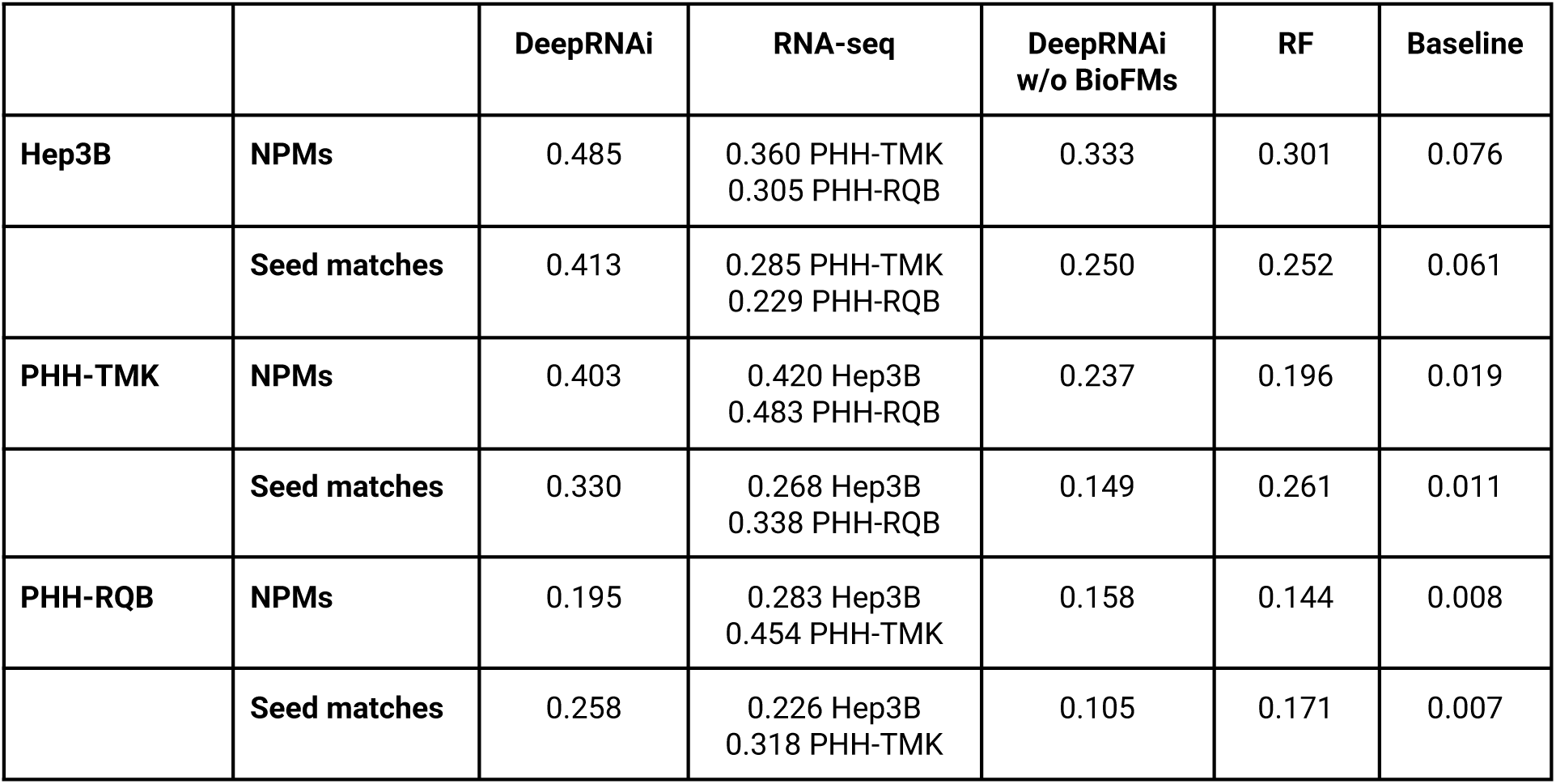
DeepRNAi accuracy on novel cell lines compared to RNA-seq cross-cell-line reproducibility. Average precision of DeepRNAi versus observed cross-cell-line RNA-seq measurements for NPMs and seed matches on 26 siRNAs screened in Hep3B and PHH derived from two different donors. Two values are listed under RNA-seq as we calculated the average precision from each of the two other cell lines.

|  |  | **DeepRNAi** | **RNA-seq** | **DeepRNAi<br>w/o BioFMs** | **RF** | **Baseline** |
|---|---|---|---|---|---|---|
| **Hep3B** | **NPMs** | 0.485 | 0.360 PHH-TMK<br>0.305 PHH-RQB | 0.333 | 0.301 | 0.076 |
|  | **Seed matches** | 0.413 | 0.285 PHH-TMK<br>0.229 PHH-RQB | 0.250 | 0.252 | 0.061 |
| **PHH-TMK** | **NPMs** | 0.403 | 0.420 Hep3B<br>0.483 PHH-RQB | 0.237 | 0.196 | 0.019 |
|  | **Seed matches** | 0.330 | 0.268 Hep3B<br>0.338 PHH-RQB | 0.149 | 0.261 | 0.011 |
| **PHH-RQB** | **NPMs** | 0.195 | 0.283 Hep3B<br>0.454 PHH-TMK | 0.158 | 0.144 | 0.008 |
|  | **Seed matches** | 0.258 | 0.226 Hep3B<br>0.318 PHH-TMK | 0.105 | 0.171 | 0.007 |

**Table S6.** Partial correlation of siRNA sequence features and RNA accessibility with observed log_2_FC knockdown in HepG2 cells, with and without DeepRNAi predictions regressed out.

|  | **On-target** |  |  |  | **NPMs** |  |  |  |
|---|---|---|---|---|---|---|---|---|
| **Approach** | Pearson corr | p-value | Pearson partial corr | Partial p-value | Pearson correlation | p-value | Pearson partial corr | Partial p-value |
| DeepRNAi | 0.80 | <10<sup>-12</sup> | - | - | 0.42 | <10<sup>-100</sup> | - | - |
| HepG2 eSHAPE | -0.21 | 0.120 | 0.23 | .0955 | -0.08 | <10<sup>-17</sup> | -0.02 | .036 |
| GC content | 0.46 | 3x10<sup>-4</sup> | 0.10 | 0.487 | 0.00 | 0.861 | 0.03 | 0.002 |
| SeedTM | 0.39 | 0.003 | -0.17 | 0.205 | 0.00 | 0.623 | 0.04 | 3x10<sup>-5</sup> |

**Table S7:** DeepRNAi and competitor performance for models trained and evaluated on the public data set (i.e. the “mini” models).

|  | **On-target** |  | **NPMs** |  | **Seed matches** |  |
|---|---|---|---|---|---|---|
| **Model** | Pearson correlation | p-value vs mini-DeepRNAi | Average Precision | p-value vs mini-DeepRNAi | Average Precision | p-value vs mini-DeepRNAi |
| mini-DeepRNAi | **0.552** | - | **0.279** | - | **0.254** | - |
| mini-DeepRNAi w/o BioFMs | 0.462 | 0.271 | 0.150 | 0.001 | 0.156 | <0.001 |
| mini-Random Forest | 0.458 | 0.322 | 0.123 | 0.001 | 0.121 | <0.001 |
| Baseline rate | - | - | 0.062 | - | 0.067 | - |

